# *REL* deregulation stands as a primary hit for AID-imprinted B-cells along the germinal center competition

**DOI:** 10.1101/2023.10.10.561773

**Authors:** Léa Prévaud, Christelle Vincent-Fabert, Tiffany Marchiol, Quentin Lemasson, Catherine Ouk, Claire Carrion, Michel Cogné, Jean Feuillard, Nathalie Faumont

**Author notes:** Corresponding author : Nathalie Faumont, UMR CRIBL CNRS7276/INSERM1262, Centre de Biologie et de Recherche en Santé, CHU et Faculté de Médecine de Limoges, Av Bernard Descottes, 87025 Limoges Cedex, France.

## Abstract

In diffuse large B-cell lymphomas (DLBCLs), gains and amplifications of the 2p15-16 region, which always encompass the *REL* gene, are mostly restricted to the germinal center (GC) B-cell DLBCL subtype (GCB-DLBCL) for which c-Rel is the pivotal Rel/NF-κB subunit. While *REL* is also known to play a key role in the GC reaction, its contribution to GCB-DLBCL transformation is still unclear. To understand the role of *REL* in the very first steps of GCB transformation, *i.e* when B-cells with deregulated *REL* are competing with other B-cells during chronic antigenic stimulation, we have created a dual-color mouse that allows to induce *REL* in a limited pool of AID- imprinted B-cells after immunization and to differentially stain AID-imprinted B-cells cells that overexpress *REL* or not. Our results demonstrate that dysregulation of *REL* at the GC B-cell stage promotes GC B-cell expansion and favors both class-switch recombination and plasma cell differentiation. Additionally, although *REL* overexpression was neutral on post-GC memory B-cell differentiation, it did confer a long-term competitive advantage allowing for GC persistence and continuous recirculation of *REL*-overexpressing B-cells. Functionally, *REL* enhanced the protection against apoptosis in the early steps of GCB differentiation. *REL*- overexpressing B-cells can my occasionally transform into in an aggressive B-cell tumor. Highlighting the role of repeated immune responses, our results confirm the role of *REL* in the germinal center reaction and provide evidence supporting the fact that genetic deregulation of c-Rel expression is most likely a primary event in the aggressive transformation of GC B-cells.

**Key points:**

- *REL* provides a long-term competitive advantage allowing for GC B-cell persistence and continuous recirculation of AID-imprinted B-cells
- AID-imprinted B-cells overexpressing *REL* can occasionally transform into aggressive B-cell lymphomas

**Explanation of the novelty:** By showing in a new dual-color mouse model that dysregulation of *REL* in a very limited pool of AID-imprinted B-cells confers a strong long-term competitive advantage in the context of repeated immune responses and may occasionally lead to transformation into an aggressive B-cell lymphoma, we provide for the first time experimental evidence supporting the fact that that *REL* is most likely a primary event in the aggressive transformation of germinal center B-cells.

## Introduction

C-Rel, the cellular equivalent of v-Rel, the oncogenic protein of Reticuloendotheliosis virus REV-T, a causative agent of lymphoid tumors in poultry, gave its name to the Rel homology domain (RHD) DNA-binding domain, that characterizes the Rel/NF-κB transcription factor family^1^. Rel/NF-κB transcription factors are composed of five sub-units, RelA, c-Rel, RelB, p50 and p52, the three formers with a transcriptional activation domain. The five Rel/NF-κB subunits associate with each other to form transcriptionally active dimers. These complexes are retained in the cytoplasm by interaction with their inhibitors, the IκΒs. Resulting from a variety of external signals, NF-κB activation consists in the proteosomal degradation of the IκΒs, and the nuclear translocation of Rel/NF-κB dimers where they regulate numerous genes critical for lymphocyte development, innate and adaptive immunity, inflammation as well as for control of both cell proliferation and cell death^2^. The canonical (or classical) NF-κB activation pathway involves nuclear translocation of RelA or c-Rel containing dimers whereas the non-canonical (alternative) pathway corresponds to nuclear translocation of RelB complexes^3^. In contrast to RelA and RelB, c-Rel does not contain a nuclear export signal and can accumulate in the nucleus upon chronic NF-κB activation^4^.

It is now very well established that constitutive NF-κB activation is the hallmark of several B-cell neoplasms, either indolent such as Waldenström Macroglobulinemia (WM), which harbors a MYD88 activating mutation in more than 90% cases, or aggressive lymphomas such as Epstein Barr virus (EBV) related lymphoproliferative disorders with expression of the EBV oncogenic protein LMP1 or diffuse large B-cell lymphomas (DLBCLs) with an activated phenotype (ABC-DLBCLs). In this latter group, NF-κB activation is genetically related to mutations in the NF-κB activation track such as those of MYD88, CD79A, CARD11 or TNFAIP3^5^.

It has only recently been recognized that the three NF-κB subunits RelA, RelB and c-Rel may have distinct and specific roles in B-cell lymphomagenesis, the latter being associated with DLBCLs with a germinal center (GC) B-cell phenotype (GCB-DLBCLs)^6–8^. In DLBCLs, 2p15-16 gains/amplifications, which include *REL*, are almost restricted to the GCB-DLBCL subtype^9,10^, being found in 15-37% of cases^11^ while it is almost never found in ABC-DLBCLs^9^. Some reports indicate that the minimal common region of 2p15-16 gains/amplifications always includes *REL*^12^. *REL* gains/amplifications, which are the first genetic aberration of the Rel/NF-κB system reported in DLBCLs^13^, are recurrently found in various B-cell cancers such as classical Hodgkin lymphoma, primary mediastinal B cell lymphoma, or chronic lymphocytic leukemia^14^. Such *REL* imbalances are also found in 20-35% of follicular lymphoma (FL) in transformation (tFL), a poor prognosis FL subtype, close to and with a similar mutational pattern than GCB-DLBCLs^11^. In contrast with other NF-κB subunits, c-Rel overexpression is very likely to be sufficient to obtain the c-Rel effects^7,15,16^. C-Rel plays a critical role in sustaining prolonged NF-κB responses in B cells and chronic BCR stimulation results in nuclear c-Rel accumulation with upregulation of pro-survival genes, whereas nuclear RelA is almost undetectable under these conditions^15^. The REL-/- mice exhibit reduced proliferation and activation of mature B cells in response to immunization associated with impaired GC formation^17^. In normal B-cells, c-Rel is a key transcription factor for GC reaction long-term maintenance^18^. In GCs, c-Rel and c-Myc would be active in the same B-cells and c-Rel would be involved in a metabolic program that would facilitate cell growth^19^. Therefore, there is robust genetic and pathophysiological evidence supporting the involvement of c-Rel in the aggressive transformation of GC B-cells.

If *REL* is a good candidate oncogene for aggressive GC B-cell transformation, it remains unclear whether it is a primary or secondary genetic event. As exemplified by FLs, a primary genetic event would occur in a very limited number of B-cells. These hit B-cells would compete with their normal counterparts for years or even decades in the context of a complex and evolving immune environment^20^. To address the question of the role of *REL* at the very first steps of GC B-cell transformation, we developed a dual-color mouse model that allows to induce c-Rel overexpression in a limited pool of AID-imprinted B-cells after immunization with a complex antigen and to differentially stain AID-imprinted B-cells cells that overexpress *REL* or not.

## Material and Methods

### Generation of mouse models (detailed in Supplemental Materials and Methods)

The conditional REL-YFP mouse model was generated for this study. The targeting vector pROSA-mREL-YFP was obtained by insertion of a 3521 pb AscI/AscI fragment containing the murine *REL* cDNA followed by an internal ribosome entry sequence (IRES) for the expression of the yellow fluorescent protein (YFP) coding sequence (the REL-YFP transgene, synthetized by GeneCust, Luxembourg), into the previously published pROSA26-1 vector^21^.

The AID.Cre^ert2^-TOM model (thereafter called AID-TOM) was obtained by crossing previously published AID.Cre^ert2^ and Ai14 mouse models^22^. AID-TOM or CD19-Cre mice^23^ and REL- YFP mice were crossed to induce the expression of the transgene in AID- imprinted or CD19pos B cells respectively: the REL-AID and REL-CD19-Cre mouse models.

### Tamoxifen induction and immunization

Immunization consisted of an intraperitoneal injection of 2.10^9^ sheep red blood cells (SRBCs) on day 0 and monthly thereafter. Tamoxifen was purchased from Gibco (Gibco Thermo Fisher Scientific France, Illkirch-Graffenstaden, France). Tamoxifen was administrated by gavage at 40 mg/kg on day 0, 2 and 4.

### Flow cytometry analysis from blood, and lymphoid organs

Blood samples were collected retro-orbitally. Spleen and lymph node cells were filtered on a 70 µm filter (Miltenyi Biotech SAS, Paris France). Peritoneal lavage was performed with three mL of phosphate-buffered saline. After red blood cell lysis, cells were labeled with fluorescent conjugated monoclonal antibodies listed in Supplemental Materials and Methods. Flow cytometry was performed with the CytoFLEX LX apparatus (Beckman Coulter France, Villepeinte, France). Results were analyzed using the Kaluza software (Beckman Coulter).

### Stimulation and Cell Culture

For proliferation analysis, splenocytes were labeled with five µM CellTraceTM Violet reagent (Invitrogen, Carlsbad, CA). Cells were then stimulated for four days with either CD40, IL4, ODN CpG or LPS as detailed in Supplemental Materials and Methods and analyzed by flow cytometry using a CytoFLEX LX (Beckman Coulter) after B220 labeling.

### Ex vivo induction of germinal center B-cell differentiation

Splenic B cells were negatively sorted using the Pan B cell isolation kit (Miltenyi Biotec). Induction of germinal center B-cell differentiation was done following the protocol of T. Nojima et al (details in Supplemental Materials and Methods).

### Statistical analysis

Statistical analyses were conducted using the GraphPad software (GraphPad Software, Boston, MA). These included Student t-tests, chi-square tests, non-parametric tests and analysis of variance (ANOVA) were performed with the GraphPad software.

## Results

### B-cells overexpressing *REL* or not following AID-induction can be differentially fate-mapped

To understand the consequences of *REL* constitutive overexpression at the germinal center B cell stage and beyond, we first generated the REL-YFP mouse model. Preceded by a STOP-Neomycin (STOP-Neo) cassette flanked by loxP sites, the murine *REL* transgene was introduced in the Rosa26 locus on chromosome 6 (Figure 1A, top panel). The *REL* transgene is followed by an IRES to allow concomitant expression of the yellow fluorescent marker YFP. In parallel, we modified the tamoxifen-regulatable AID.Cre-ert2 mouse model of Dogan et al.^22^ to introduce a sequence encoding the red tdTomato marker preceded by a STOP cassette flanked by loxP sites (Figure 1A). Mice were wisely selected to retain those in which a crossing-over event placed the flox_tdTomato transgene in cis to AID.Cre-ert2 allele at the Rosa26 locus: the AID-TOM model (figure 1A, lower panel). Breeding REL-YFP with AID-TOM mice resulted in the dual-color REL-AID model. In this model, the two tdTomato and REL-YFP transgenes are both AID-inducible and tamoxifen-regulatable.

**Figure 1:**
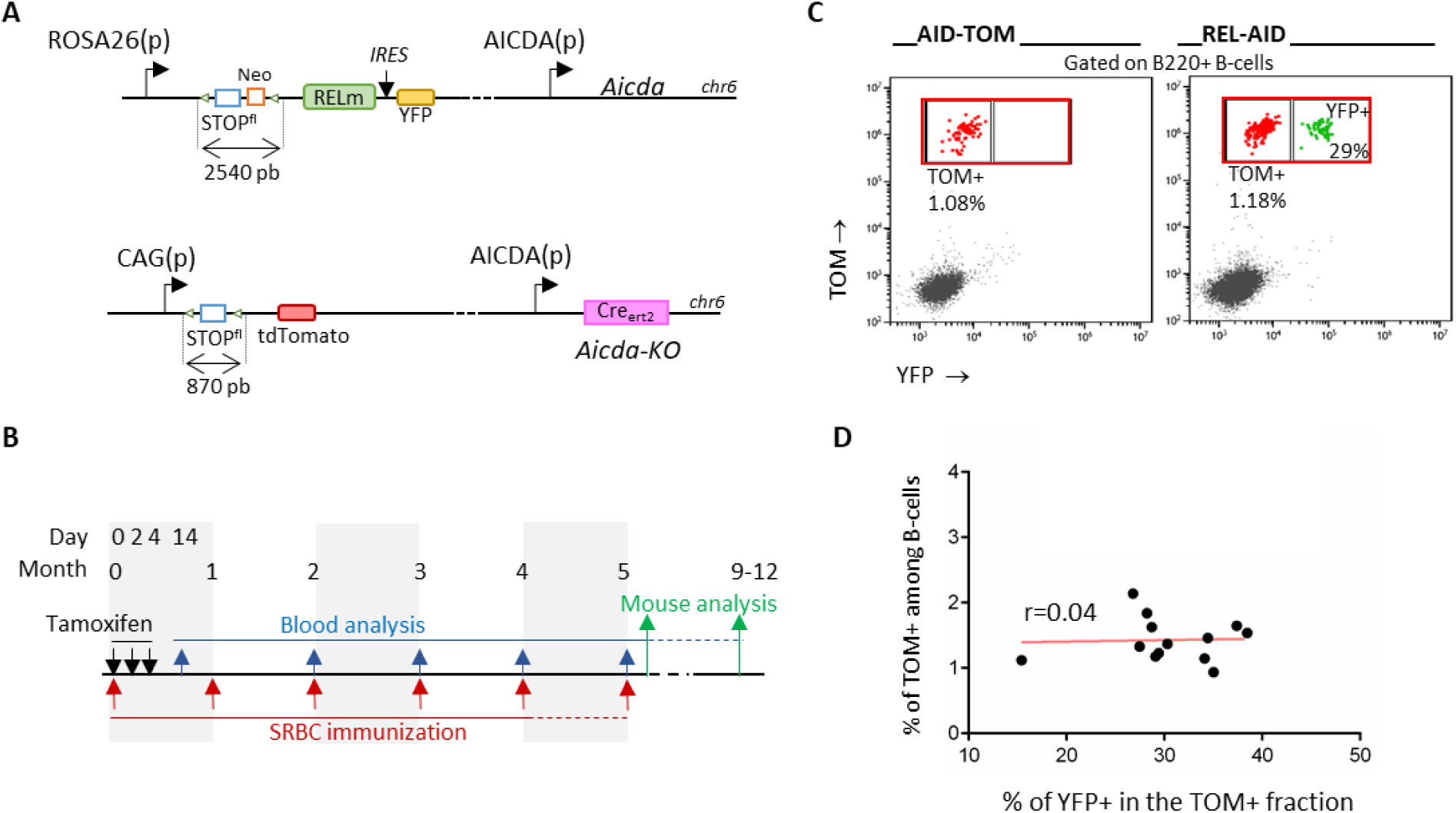
Characterization of the REL-AID mouse model. A: Schematic representation of the REL-IRES-Yfp and tdTomato inserts into the Rosa26 locus: the *REL* sequence was placed in frame with the Internal Ribosomal Entry Site (IRES) and the coding sequence for Yellow Fluorescent Protein (YFP). B: Schematic diagram of the immunization protocol for analysis either shortly or long time after the end of immunizations. Blood samples were collected on D14 and then each month, always before immunization recall throughout the immunization and post-immunization period. C: Example of a flow cytometry biparametric YFP and tdTomato (TOM) histogram gated on live B220pos B-cells. TOM+/YFP- and TOM+/YFP+ B-cells are colored in red and green respectively. Circulating blood cells were collected from a AID-TOM (left) or a REL- AID (right) mouse one month after immunization and tamoxifen gavage. Percentages of fluorescent cells are shown in each graph. D: Relationship between the percentage of TOM+/YFP+ among total TOM+ B-cells (x-axis) and that of TOM+ B-cells among total B220pos B-cells (y-axis) in REL-AID mice. The correlation curve is shown in red. The value of the correlation coefficient r is shown in the graph.

To induce a very limited number of B-cells in the context of complex repeated immune responses, our protocol consisted of an initial tamoxifen Cre-ert2 induction together with chronic immunization with a complex T-dependent antigen, in this case sheep red blood cells (SRBC) (Figure 1B). Using AID-TOM mice as a control, we intraperitoneally injected SRBCs on day 0 (D0) together with tamoxifen gavage on D0, D2 and D4 (Figure 1B), which would favor initial germinal center formation. For long term GC maintenance, monthly SRBC immunization recalls were conducted for four or five months. At month 5 (M5), mice were either sacrificed shortly after chronic immunization or monitored for six to eight months before being sacrificed in order to study the persistence of TOM+/YFP+ B cells long time after the end of immunization (Figure 1B).

As shown in figure 1C, circulating TOM+ B-cells could be detected in both AID-TOM and REL-AID mice at D14, accounting for 0.79% - 2.14% of total circulating B-cells, with no significant differences between the two mouse strains. As the Cre efficiency is inversely dependent on the size of the cassette to be excised between the loxP sites, not all B-cells that had deleted the 870 bp long STOP cassette of the AID.Cre-ert2 allele had also eliminated the 2540 bp long STOP-Neo cassette of the wild-type AID allele. At D14, 28.5% +/- 2.4% of TOM+ circulating B-cells were also YFP+ (figure 1D). Therefore, thanks to the YFP and tdTomato reporters, this dual-color model allows for differentiated fate-mapping of TOM+/YFP+ (RELpos) and TOM+/YFP- (RELneg) B cells upon tamoxifen administration after AID-imprinting (likely to be due to germinal center passage) within the same animal.

### Long-term competitive advantage of *REL* overexpressing B-cells when compared to their *REL* negative counterpart

After initial tamoxifen induction of Cre-ert2 activity in AID-TOM control mice, the levels of circulating TOM+ B-cells moderately decreased despite SRBC immunization recall. Then, TOM+ B-cell percentages remained stable as long as mice were repeatedly immunized, implying an equilibrium or a stationary state between blood entry and exit of TOM+ B-cells. This would indicate a continuous recirculation and/or production of TOM+ B cells in secondary lymphoid organs. Subsequently, TOM+ B-cells gradually diminished to near-zero levels as immunization recalls ended, revealing the reliance of TOM+ B-cell production. This indicates a continuous recirculation and/or production of TOM+ B cells in secondary lymphoid organs.

Then, TOM+ B-cells progressively reached levels close to zero as immunization recalls stopped, which shows the dependency on the immune response. Contrary to AID-TOM mice, REL-AID mice exhibited stable levels of circulating TOM+ B-cells not only during the immunization period but also for up to two months after cessation of immunization before declining (Figure 2A). This shows that *REL* overexpression led to an increase in production (blood entry increase) and/or survival (blood exit decrease) of TOM+ B-cells. The extent of this increase was also influenced by the persistence of the immune response.

**Figure 2:**
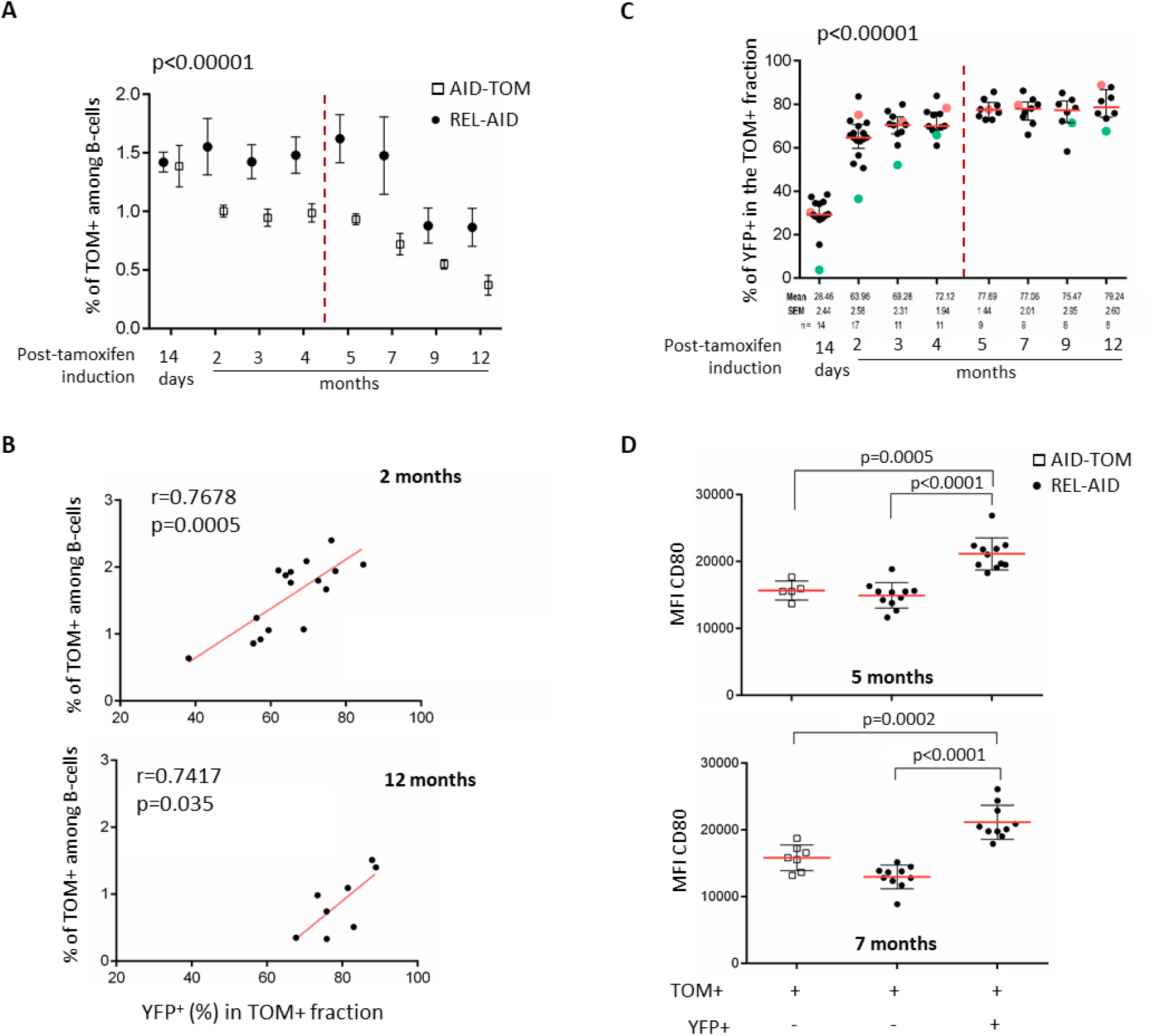
Relationships between the kinetics of circulating TOM+ and TOM+/YFP+ with CD80 expression. A: Box plot of the percentage of circulating TOM+ cells in AID-TOM and REL-AID mice over time. The dashed red line indicates the end of immunization recall. The p-values of a two-way ANOVA for REL effect is shown. B: Relationship between the percentage of TOM+/YFP+ among total TOM+ B-cells (x-axis) and that of TOM+ B-cells among total B220pos B-cells (y-axis) in REL-AID mice at month 2 (top graph) and month 12 (bottom graph). The correlation curve is shown in red. The values of the Pearson correlation coefficient r and of its p-values are given in each graph. C: Distribution of the percentage of TOM+/YFP+ B-cells among total TOM+B-cells over time in REL-AID mice. The red dot represents the mouse with the highest percentage and green dot the mouse with the lowest percentage. The p-value of the one-way ANOVA is shown within the graph. D: CD80 expression levels in TOM+/YFP- from either AID-TOM and REL-AID as well as TOM+/YFP+ mice. Mann-Whitney test p-values are shown

As shown in figure 2B and 2C, the increased production and/or survival of TOM+ B-cells in REL-AID mice was due to TOM+/YFP+ double positive, i.e to *REL* overexpressing B-cells. Indeed, percentages of TOM+ cells among total circulating B-cells were strongly correlated with those of YFP+ cells within the TOM+ B-cell fraction at M2 and M12 (Figure 2B), which is in contrast with the result at D14 (Figure 1D). Figure 2C shows the progressive but rapid, consistent and persistent accumulation of YFP+ B-cells within the TOM+ B-cell compartment, even when immunization recalls ceased. Compared to TOM+ B-cells from AID-TOM mice or TOM+/YFP- B-cells from REL-AID mice, TOM+/YFP+ B-cells over-expressed CD80 (Figure 2D), a marker of NF-κB that is known to be express on a subset of memory B-cells^24^ and to be increased upon c-Rel activation^25^.

Collectively, these results first demonstrate that recirculation of AID imprinted B-cells is highly dependent on chronic immunization. They also strongly support a significant long-term competitive advantage of B-cells that have benefited from a *REL* overexpression event when compared to their *REL* negative counterpart.

### Expansion and persistence of REL-overexpressing B-cells in secondary lymphoid organs

Shortly after immunization, the proportion of TOM+ B-cells was increased in the peritoneum, i.e at the SRBC injection site, when compared to the spleen or lymph nodes in AID-TOM control mice (Figure 3B), again indicating the role of antigen stimulation in promoting AID- imprinted B-cells. This phenomenon was markedly increased in REL-AID mice, as TOM+ B- cells ranged from 10% to 50% of total B-cells in the peritoneum (Mann-Whitney test, p=0.057). Although heterogeneous, levels of TOM+ B cells were also increased in spleen and lymph nodes of REL-AID mice, and most of these TOM+ B-cells were also YFP+. Spleen histology was similar in both control AID-TOM and REL-AID mice with large and somewhat coalescing lymphoid nodules and expanded B-cell areas (Figure 3C) as well as numerous plasma cell sheets in the red pulp (supplemental figure S1). However, one REL-AID mouse of this cohort, marked with an arrow in Figure 3, developed a diffuse large B-cell lymphoma (supplemental figure S2). In this mouse, almost all TOM+ B-cells were also YFP+, i.e were overexpressing *REL* (Figure 3B, lower panel).

**Figure 3:**
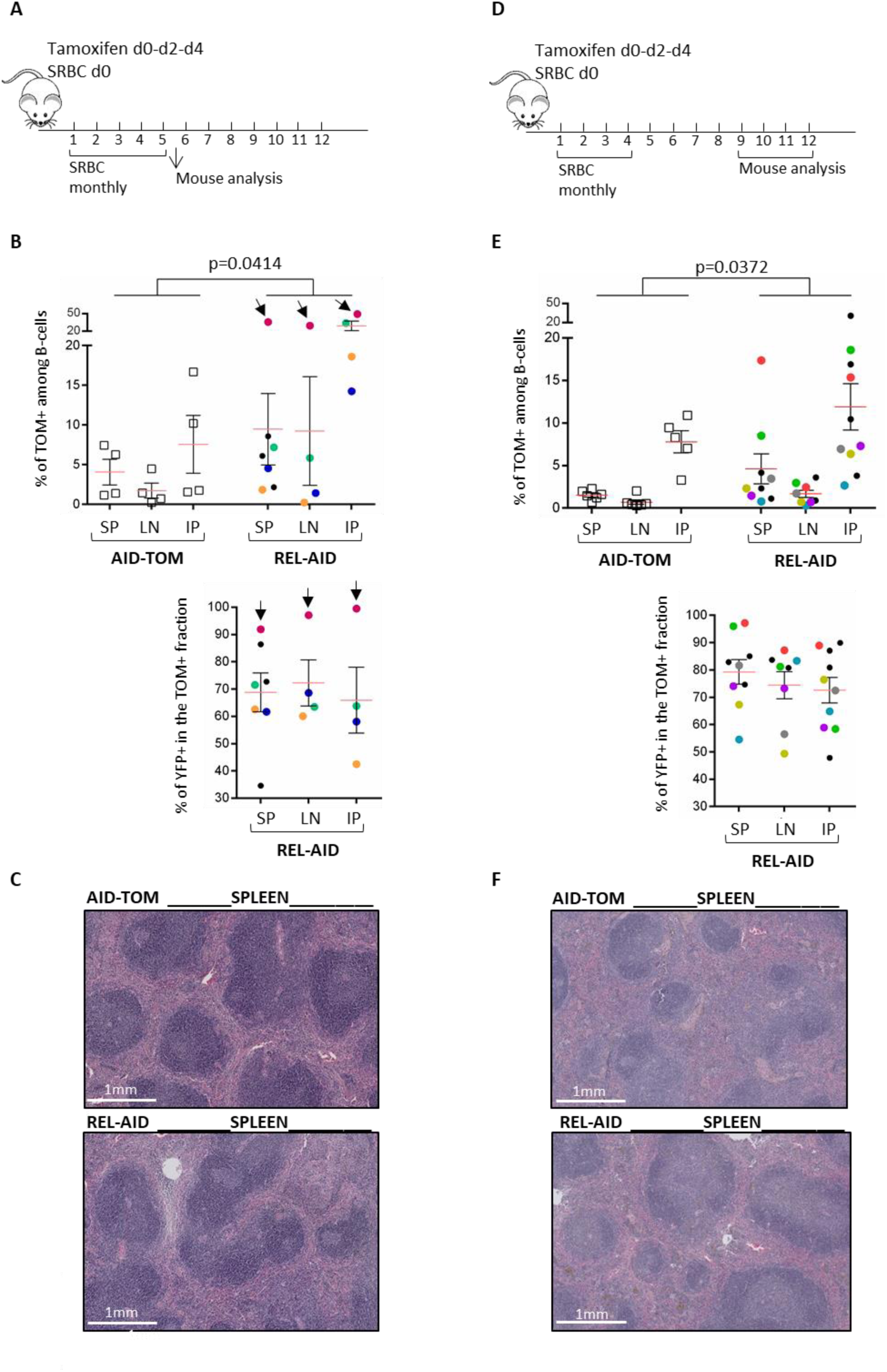
Relationship between the levels of TOM+ and TOM+/YFP+ B-cells in secondary lymphoid organs. A and D: Schematic representation of the immunization protocol for analysis either shortly (Figure 3A) or long time (Figure 3D) after the end of immunization. B and E: Analysis of TOM+ and TOM+/YFP+ B-cells in secondary lymphoid organs shortly (Figure 3B) or long time (Figure 3E) after the end of immunization for AID-TOM control (left) and REL-AID mice (right) in the spleen (SP), in lymph nodes (LN) and in the peritoneum (IP). Top: percentage of TOM+ B-cells among total live B-cells. Bottom: percentage of TOM+/YFP+ B-cells among TOM+ B-cells. In Figure 3B, the case with a diffuse aggressive B-cell lymphoma is indicated by an arrow. P-values are shown on the top of the graph. C and E: Histological aspect of the spleen shortly (Figure 3C) or long time (Figure 3F) after the end of immunization for AID-TOM control (top) and REL-AID mice (bottom).

Long time after immunization, while persisting in the peritoneum, TOM+ B-cells almost disappeared from both spleen and lymph nodes of AID-TOM mice. In contrast, TOM+ B-cells persisted, albeit at lower levels and more heterogeneously than immediately after immunization in both spleen and lymph nodes and remained higher in the peritoneum of REL-AID mice. Again, most TOM+ B-cells from REL-AID mice were also YFP+ (Figure 3E). Reflecting resting spleens, spleen histology showed small lymphoid nodules with increased red pulp areas in 8/9 (89%) control AID-TOM. In contrast REL-AID mice tended to retain large lymphoid nodules with increased B-cell areas in 12/14 (86%) cases (Khi2 test, p=7.10^-4^ and Figure 3F). Taken together, these results show that *REL* promotes the expansion and persistence of AID- imprinted B-cells in the secondary lymphoid organs.

### REL-overexpressing germinal center B-cells display a competitive advantage and are prone to immunoglobulin switching

The fluorescent markers TOM and YFP were used to define the splenic B-cell composition of RELpos (TOM+/YFP+) and RELneg (TOM+/YFP-) B-cells in REL-AID mice shortly after immunization (Figure 4A). CD95pos CD38low GC B-cells were significantly increased in TOM+ B-cells of REL-AID mice when compared to their AID-TOM control (Figure 4B). This GC B-cell fraction was specifically enriched in RELpos B-cells, reaching almost 90% of total TOM+ B-cells, even in mice with a rather low percentage of YFP+ cells among total TOM+ B-cells (Figure 4C). Indeed, the GL7 positive GC B-cell fraction was found almost exclusively in RELpos B-cells (Figure 4D). Interestingly, the increase in GC B-cells in the TOM+ fraction persisted a long time after immunization was stopped in REL-AID mice, whereas these GC TOM+ B-cells almost disappeared in AID-TOM controls (Supplemental Figure S3).

**Figure 4:**
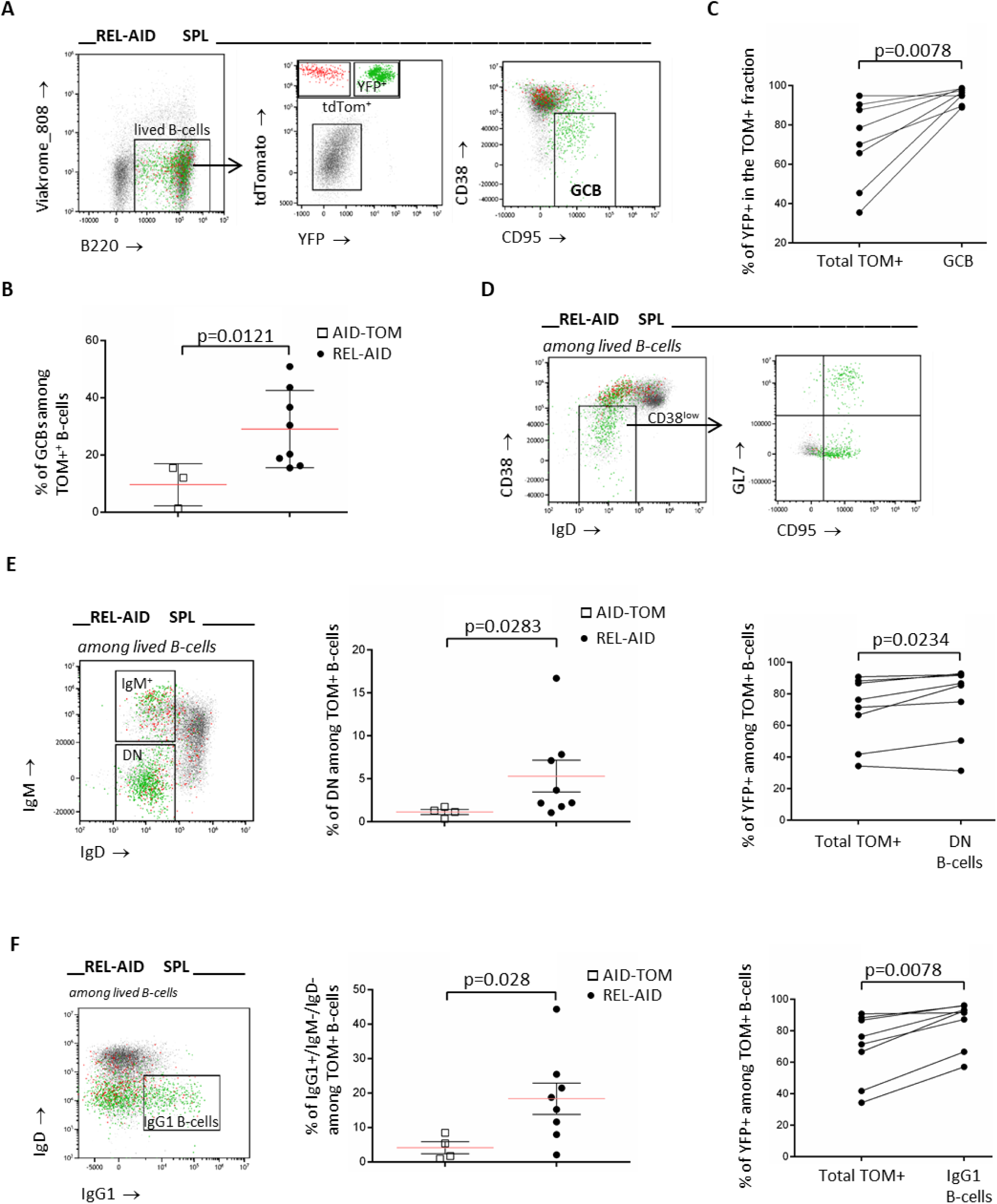
Analysis of splenic germinal center and IgG1 switched B-cells shortly after immunization. A: Gating strategy for gating TOM+/YFP- (colored in red) and TOM+/YFP+ (colored in green) as well as germinal center B-cells (GCBs) from total live B-cells. B: Percentage of GCBs among TOM+ B-cells in AID-TOM control and REL-AID mice. The p-value of the Mann-Whitney test is shown. C: Percentage of TOM+/YFP+ B-cells in the total TOM+ (left) or in the GCB (right) fraction in REL-AID mice. Each line connects the points for one mouse. The p-value of the Wilcoxon test is shown. D: Example of gating on the CD38low/IgDneg B-cell fraction (left) showing that almost all B-cells with a CD95+/GL7high GCB phenotype (right) are colored in green, corresponding to TOM+/YFP+ B-cell cells. E: Analysis of IgMneg/IgDneg double negative (DN) B-cells: left: example of gating on DN B-cells; middle: percentages of DN cells among total TOM+ B-cells in AID-TOM and REL-AID mice; right: percentages of TOM+YFP+ B-cells among TOM+ B-cells in the total TOM+ and in the DN B-cell fraction in REL-AID mice (each line connects the points for one mouse). The p-values of the Mann-Whitney (middle) and Wilcoxon test (right) are shown. F: Analysis of IgG1 switched B-cells: left: example of gating on IgG1+ B-cells; middle: percentages of IgG1+ DN B-cells among total TOM+ B-cells in AID-TOM and REL-AID mice; right: percentages of TOM+YFP+ B-cells among TOM+ B-cells in the total TOM+ and in the IgG1+ DN B-cell fraction in REL-AID mice (each line connects the points for one mouse). The p-values of the Mann-Whitney (middle) and Wilcoxon test (right) are shown.

RELpos B-cell enrichment was not found for IgMpos IgDneg B-cells (Figure 4E and Supplementary Figure S4). Including switched B-cells, the IgM/IgD double negative B-cell subset was moderately increased in REL-AID mice when compared to TOM-AID controls. The enrichment in RELpos B-cells in this subset was moderate although significant (Figure 4E). But IgM/IgD double negative B-cells also contain GC B-cells that have downregulated their BCR or have a nonfunctional or damaged BCR. Thus, we also looked at the expression of the IgG1 isotype. As shown in figure 4F, although heterogeneous, percentages of IgG1 switched B-cells were increased in REL-AID mice and these IgG1 B-cells were enriched in RELpos B- cells.

### REL-overexpressing B-cells are prone to plasma cell but not to memory B-cell differentiation

Targeting plasma cells (PCs) with the CD138 marker revealed that PC differentiation was rather heterogeneous in REL-AID mice, although not significantly increased when compared to TOM-AID controls. But, almost all PCs from REL-AID mice were derived from the RELpos B-cell fraction (figure 5A).

**Figure 5:**
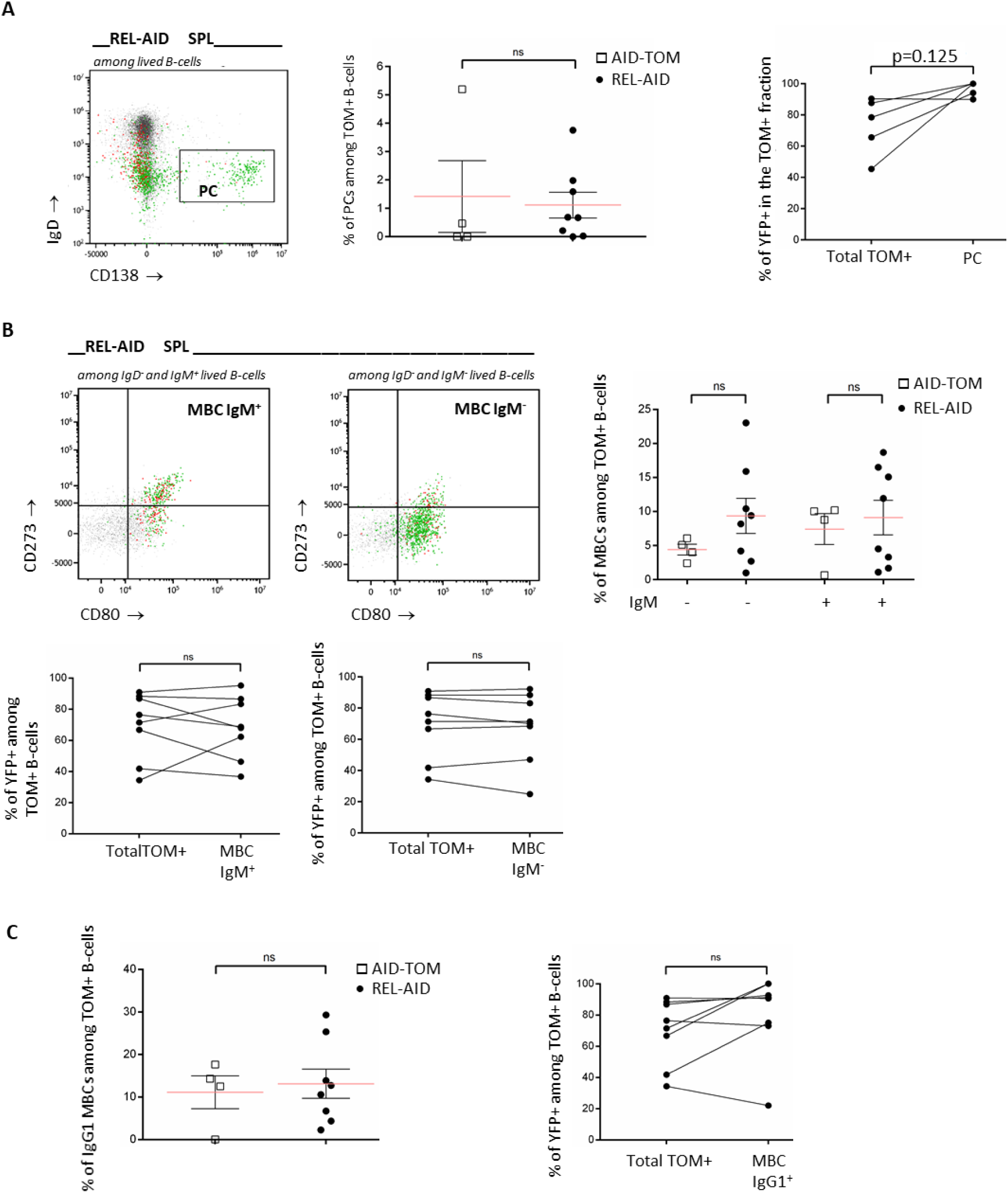
Analysis of splenic plasma cells and memory B-cells shortly after immunization. A: Analysis of plasma cells (PCs): left: example of gating on PCs; middle: percentages of PCs cells among total TOM+ B-cells in AID-TOM and REL-AID mice; right: percentages of TOM+YFP+ B-cells among TOM+ B-cells in the total TOM+ and in the PC fraction in REL- AID mice (each line connects the points for one mouse). B: Analysis of memory B-cells (MBCs): top left: example of CD80/CD273 biparametric histogram for IgD-/IgM+ (IgM+) and IgD-/IgM- B-cells. MBCs are CD80+/CD273+; top right: percentages of IgM- and IgM+ MBCs B-cells among total TOM+ B-cells in AID-TOM and REL-AID mice; bottom: percentages of TOM+YFP+ B-cells among TOM+ B-cells in the total TOM+ and in the IgM+ (left) and in the IgM- (right) B-cell fractions in REL-AID mice (each line connects the points for one mouse). C: Analysis of IgG1 switched MBCs: left: percentage of IgG1+ MBCs among total TOM+ B-cells in AID-TOM and REL-AID mice; right: percentage of TOM+YFP+ B-cells among TOM+ B-cells in the total TOM+ and in the IgG1+ MBC fraction in REL-AID mice (each line connects the points for one mouse). ns: non-significant

Memory B-cells (MBCs) were defined by increased expression of the CD273 marker ^24^. Both IgM+ and IgM- MBCs were detected with this marker (Figure 5B). Levels of both IgM+ and IgM- MBCs were not significantly increased in REL-AID mice, although more heterogeneous than in AID-TOM mice. The percentages of RELpos B-cells in both MBC subsets was similar to that of RELneg B-cells (Figure 5B), indicating that *REL* was neutral on post GC MBC differentiation. Indeed, neither an increase of IgG1 MBCs nor an enrichment of RELpos B-cell in this fraction was found in REL-AID mice.

### CD40 stimulation increased the ex vivo proliferation of B-cells overexpressing REL

In parallel to REL-AID mice and by crossing REL-YFP with CD19-Cre mice, we also generated a REL-CD19-Cre mouse model. As expected, more than 80% of REL-CD19-Cre B-cells were YFP positive (Figures 6 and 7), while CD19 negative cells were not (not shown). These mice did not show any phenotype after 18 month follow-.up. However, we could use this model to functionally analyze the effects of *REL* on whole CD19pos B-cells. As exemplified in Figure 6A, cell proliferation was examined by dilution of the Cell-Trace violet (CTV) marker over successive mitoses. The percentage of YFP+ B-cells remained stable over the course of the ex-vivo experiment (Figure 6A). The results presented in Figure 6B show that *REL* overexpression alone did not play significantly on ex-vivo B-cell proliferation. Response to signals targeting TLR9 or TLR4 could be increased in REL-CD19-Cre B-cells, but the differences were not statistically significant when compared to controls (Figure 6B). CD40 signal appeared to weakly but consistently and significantly increase REL-CD19-Cre B-cell proliferation. REL-CD19-Cre splenic B-cells were as responsive to IL4 alone or in combination with CD40 as CD19-Cre control B-cells. Thus, at the global B-cell level, the effects of *REL* on proliferation appeared to be enhanced by CD40 stimulation.

**Figure 6.**
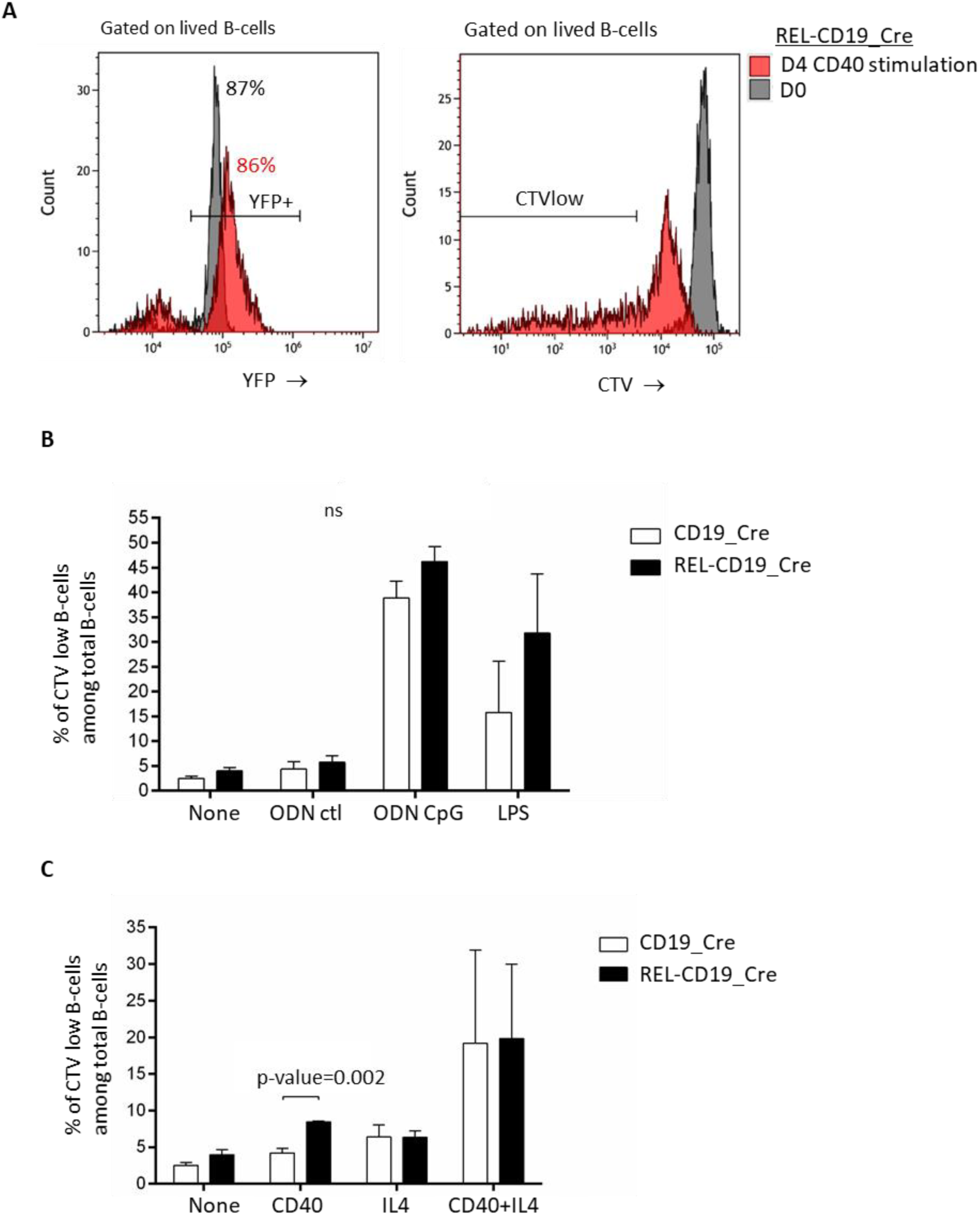
Ex-vivo proliferation assay for B-cells from CD19-Cre and REL-CD19-Cre mice. A : Example of histograms for YFP (left) and Cell Trace violet (CTV, right) at D0 (grey) and D4 (red) of ex-vivo CD40 stimulation for splenic B cells from a REL-CD19-Cre mouse. B and C: Histograms showing the percentage of CTV low B-cells for CD19-Cre and REL-CD19-Cre mice in response to ODN CpG and LPS compared to no stimulation and to a control ODN (ODN ctl) (panel B) or in response to CD40, IL4, both or no stimulation (panel C).

**Figure 7.**
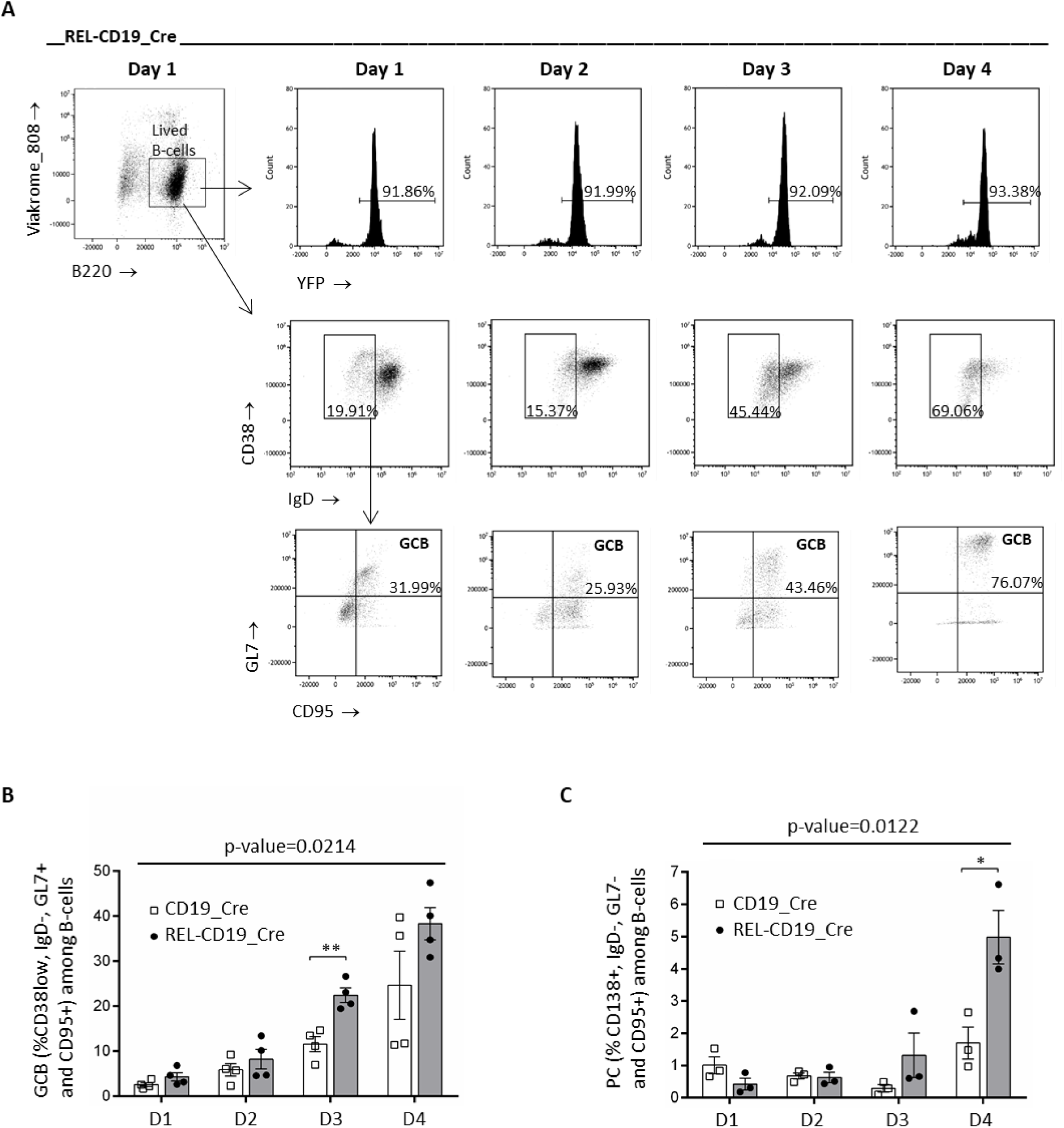
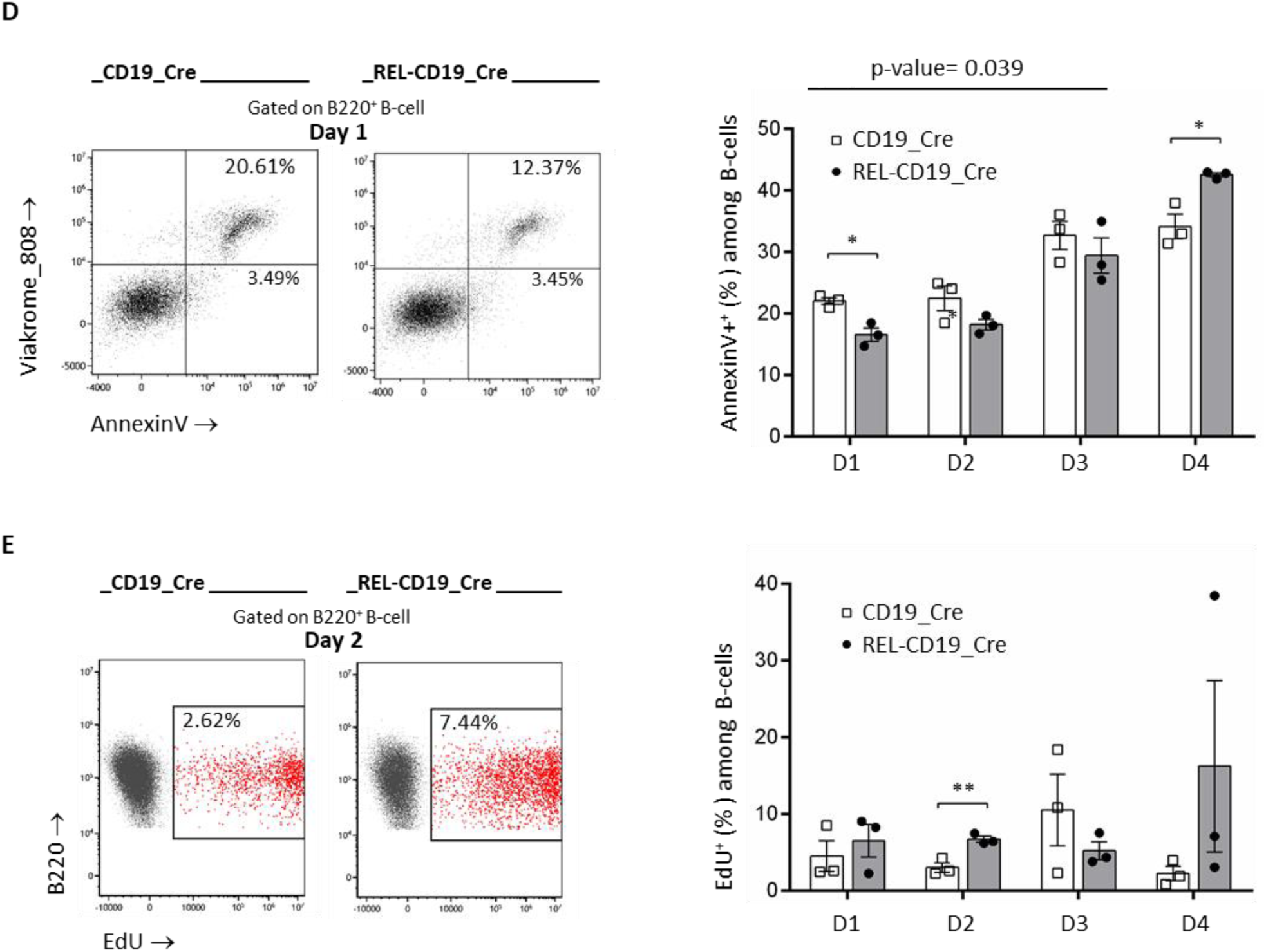
Ex-vivo germinal center B-cell and plasma cell differentiation for splenic B-cells from CD19-Cre and REL-CD19-Cre mice. A: Example of histograms for YFP, IgD/CD38 and CD95/GL during the time course of the GC B-cell differentiation for a REL-CD19-Cre mouse; top: YFP fluorescence histograms of live B-cells from Day 1 to Day 4 showing the stability of the transgene expression; middle: IgD/CD38 biparametric histograms showing the increase in the percentage of IgDneg/CD38pos or low B-cells over time; bottom: CD95/GL7 biparametric histograms showing the increase in the percentage of B-cell with a CD95high/GL7high GCB B-cell phenotype among IgDneg/CD38pos or low B-cells. B: Percentages of IgDneg/CD38pos or low/ CD95high/GL7high germinal center B-cells among total B-cells from Day 1 to Day 4 C: Percentages of plasma cells (PCs) among total B-cells from Day 1 to Day 4. D: Example of Viakrome_808/Annexin V biparametric histogram for a CD19-Cre and a REL-CD19-Cre mouse with percentages in each right quadrant (left) and percentages of Annexin V positive B-cells during the time course of the GC B-cell differentiation (right). E: Example of EdU/B220 biparametric histogram for a CD19-Cre and a REL-CD19- Cre mouse and percentages of EdU positive B-cells during the time course of the GC B-cell differentiation (right). p-values for ANOVA are shown when significant. Multiple t-test with Holm-Sidak correction were also performed (*: p<0.05; **: p<0.01)

### Initially protecting cells from apoptosis, *REL* increased ex-vivo germinal center B-cells differentiation and plasma cell production

In a next step we used the REL-CD19-Cre model to investigate germinal center B-cell differentiation *ex-vivo* according to the protocol developed by Nojima et al.^26^. Purified B- splenocytes were induced for GC B-cell differentiation by contact with CD40L and BAFF- secreting fibroblasts in the presence of IL4 for four days. As shown in Figure 7A, the percentage of YFP+ REL-CD19-Cre B-cells remained constant over time, while IgD expression was lost, CD38 surface levels tended to decrease and both GL7 and CD95 GC-B-cell markers were progressively acquired, reaching a maximum at D4. Compared to their CD19-Cre control, the percentage of REL-CD19-Cre B-cells differentiating into GC B-cells was markedly increased, with the most significant difference found at D3 (Figure 7B). Regarding post-GC differentiation, the percentage of memory B-cells, either IgM positive or IgG1 switched, was similar in both CD19-Cre control and REL-CD19-Cre mice (not shown). In contrast, plasma cell production was increased in REL-CD19-Cre mice at D3 and D4, although at low percentages (figure 7C).

We then looked at apoptosis and proliferation of ex-vivo GC-differentiating B-cells. As shown in Figure 7D, the percentage of annexin V positive REL-CD19-Cre B-cells was first decreased at D1 and D2 and then increased as the cells differentiated into GCBs, exceeding those of control cells at D4. As assessed by EdU incorporation, proliferation of GC-differentiating B-cells was rather similar between control and REL-CD19-Cre mice during the first three days and increased very heterogeneously in the latter at D4 (Figure 7E).

Altogether, these results show that *REL* overexpression in B-cell favored ex-vivo GC B-cell differentiation and plasma cell production. The effects of *REL* on proliferation seemed marginal, while those on apoptosis were more consistent for the first three days of GC B-cell differentiation.

## Discussion

Questioning the role of c-Rel in first steps of GC B-cell transformation, we present for the first time a dual-color mouse model, so-called REL-AID, in which a genetic event responsible for permanent REL overexpression is transiently induced in only a small fraction of the pool of B-cells irreversibly marked by AID imprinting, allowing color tracking of REL-overexpressing B-cells distinct from their REL-negative counterparts. Results clearly show that dysregulation of c-Rel in GC B-cells promotes GC B-cell expansion, favors CSR and PC differentiation and confers a long-term competitive advantage.

Recently, a mouse model with c-Rel overexpression in B-cells was reported by Kobel- Hasslacher et al.^16^. In this model, the *REL* transgene was under the dependence of a strong CAG promoter. *REL* overexpression was obtained in either all CD19pos or all switch-engaged GC B-cells, after breeding with CD19-Cre or Cγ1Cre mice^23,27^. The phenotype of these mice demonstrated a positive effect of c-Rel on GC B-cell expansion, resulting in increased PC differentiation with production of autoantibodies. In comparison, we could not detect any phenotype of our REL-CD19-Cre mice. Only ex-vivo functional studies on purified B-cells revealed that c-Rel could promote GC-B cell differentiation and PC production by initially protecting GC engaged B-cell from apoptosis. Our *REL* transgene was under the control of the Rosa26 promoter, which is known to be 10 times much weaker than the CAG promoter^28^. The absence of a *REL* phenotype in REL-CD19-Cre mice may very well be due to rather low levels of *REL* overexpression. Proliferation experiments showed a weak but significant c-Rel synergy only in presence of CD40. Like BAFF, CD40 is a potent activator of the alternative NF-κB activation pathway^3^. BAFF is essential for ex-vivo GC B-cell differentiation^26^. Invalidation of BAFF and BAFF receptors results in the rapid decline of germinal centers after antigenic stimulation^29^. Together with RelB, both BAFF and CD40 are also able to induce c-Rel activation^30,31^. Overall, our results raise the question of the synergy of c-Rel and RelB in the c-Rel-dependent expansion of GC B-cells.

Since induction of *REL* in all CD19pos B-cell would not reflect the true physiopathology of REL-associated B-cell transformation, we developed the dual-color REL-AID model in an attempt to get closer to what is most likely to occur in the true life of an emerging tumoral B-cell that has recently been hit by an initial genetically transforming event such as *REL* gains in GCs. *REL* overexpression in GCB DLBCLs is directly correlated with *REL* gains but is rather modest, being about 1.5 fold-change in most cases^7^. Therefore, we first chose to keep our *REL* transgene under the control of the native Rosa26 promoter to mimic a weak but continuous deregulation of REL. Second, Cre-ert2 expression was dependent on AID regulation to mainly target the GC B-cell stage. Third, Cre-ert2 activity was very transiently induced by tamoxifen gavage together with immunization with a complex antigen. Fourth, the few RELpos B-cells generated after tamoxifen gavage remained in competition with all other *REL* unhit B-cells for a long time in the context of a complex and chronic immune response. This competition would begin within GCs and would continue later in post-GC differentiation stages. To find that the small pool of RELpos AID-imprinted B-cells was favored *in vivo* for GC reaction and then for CSR and PC differentiation is in agreement with Kobel-Hasslacher et al.^16^. Moreover, the results also show a clear competitive advantage in the long-term maintenance and recirculation upon and after chronic immunization for RELpos B-cells. Because *REL* was neutral on MBC differentiation in secondary lymphoid organs, this *REL* effect on the recirculating B-cells would occur later, favoring long-term survival of post-GC recirculating MBCs. In addition to highlighting the role of chronic antigenic stimulation, our results also indicate that part of this competitive advantage may very well be related to the long-term persistence of RELpos GC B-cell production in secondary lymphoid organs.

Most B-cell transformation scenarios imply a primary event that allows escape from homeostasis and then accumulation of mutations over-time until clinical emergence of the B-cell tumor. Resulting in deregulation of Bcl2, the primary transforming event in FLs is the t(14;18)(q32;q21) translocation. These t(14;18)pos B-cells can be detected at a very low frequency in most adults. However, the vast majority of these t(14;18)pos B-cells will never transform. Clonal FL lymphomagenesis is due to the accumulation of secondary mutations such as those in CREBBP, TNFSRF14, EZH2 or KMT2D over decades^20^. These secondary mutations are also commonly found in GCB DLBCLs with the *REL* signature ^7^. By showing that the weak but continuous deregulation of *REL* favors emergence of a B-cell that will be positively selected and expanded in GCs and will have a strong long-term competitive advantage in the context of repeated immune responses, our results provide evidence for *REL* gains/amplification as a primary genetic event in the aggressive transformation of GC B cells. Consistently, and although it was only one case, one REL/AID mouse in our cohort developed an aggressive B-cell tumor with almost 100% RELpos B-cells.

Collectively, our findings suggest for the first time that genetic deregulation of c-Rel expression is most likely a primary event in the aggressive transformation of GC B-cells and raise the question of *REL*-associated secondary events for GC B-cell transformation.

## Supporting information

Legends for supplementary figures

