## Supplementary material for "*REL* deregulation stands as a primary hit for AID-imprinted B-cells along the germinal center competition": Legends for supplementary figures

**Supplementary figure 1: Immunoglobulin immunostaining of a 5µm spleen section from an AID-TOM control (top) or a REL-AID (bottom) mouse shortly after the end of immunization. A** high magnification image of a plasma cell sheet (white rectangle) is superimposed on the top right of the low magnification image. The magnification scale is indicated at the bottom of each image.

**Supplementary Figure 2: Hematein-Eosin staining of a 5µm spleen section of REL-AID mouse #7635, which developed an aggressive B-cell tumor with almost 100% TOM+/YFP+ positive B-cells.** The high magnification image of tumor cells is superimposed on the top right of the low magnification image. The magnification scale is shown at the bottom of each image.

**Supplementary figure 3: Percentage of GCBs among TOM+ B-cells in AID-TOM control and REL-AID mice long time after the end of immunization.**

**Supplementary figure 4: Percentage of TOM+/YFP+ B-cells in the total TOM+ (left) or in the IgM+/IgD- (right) fraction in REL-AID mice long time after the end of immunization. Each line connects the points for one mouse.**
