## Supplementary Figures for "*REL* deregulation stands as a primary hit for AID-imprinted B-cells along the germinal center competition"

**Supplementary figure S1. PREVAUD *et al.***

AID-TOM C3 = 863

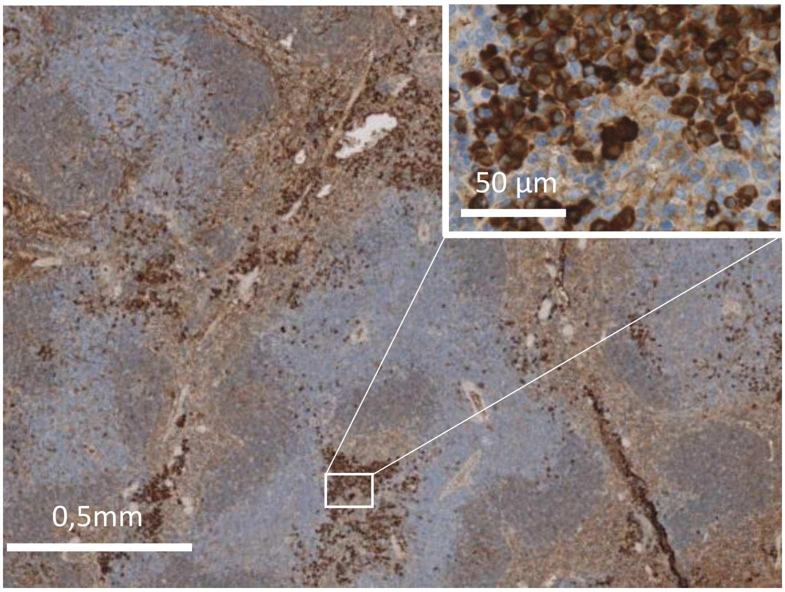


REL C3 = 851

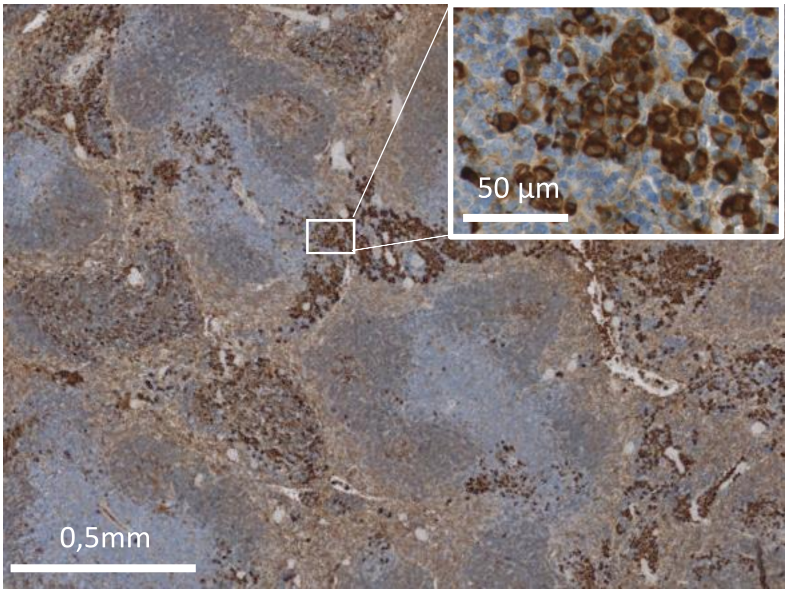

**Supplementary figure S2. PREVAUD *et al.***

AID-REL C3 = 7635

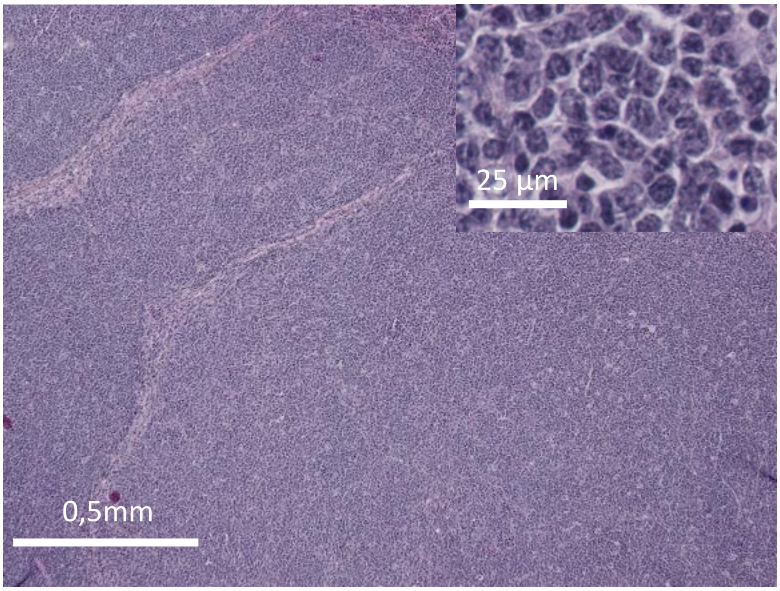

**Figure S3. PREVAUD *et al.***

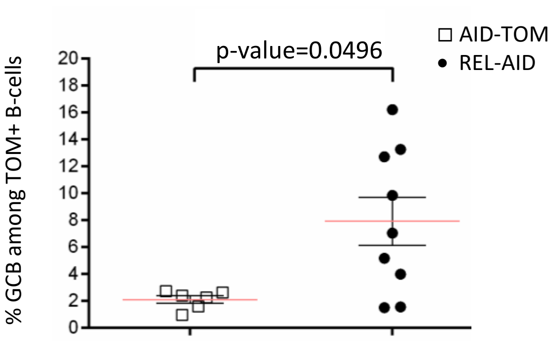

**Figure S4. PREVAUD *et al.***

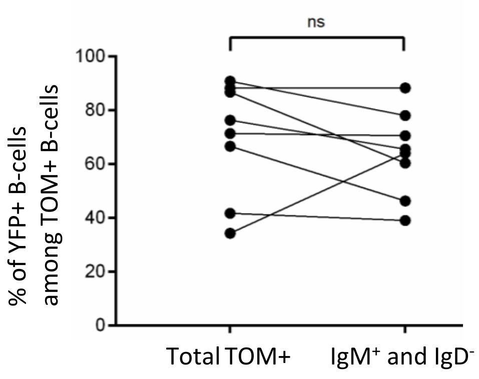
