## Supplementary Materials and Methods for "*REL* deregulation stands as a primary hit for AID-imprinted B-cells along the germinal center competition"

#### The sequence of the mREL-YFP insert

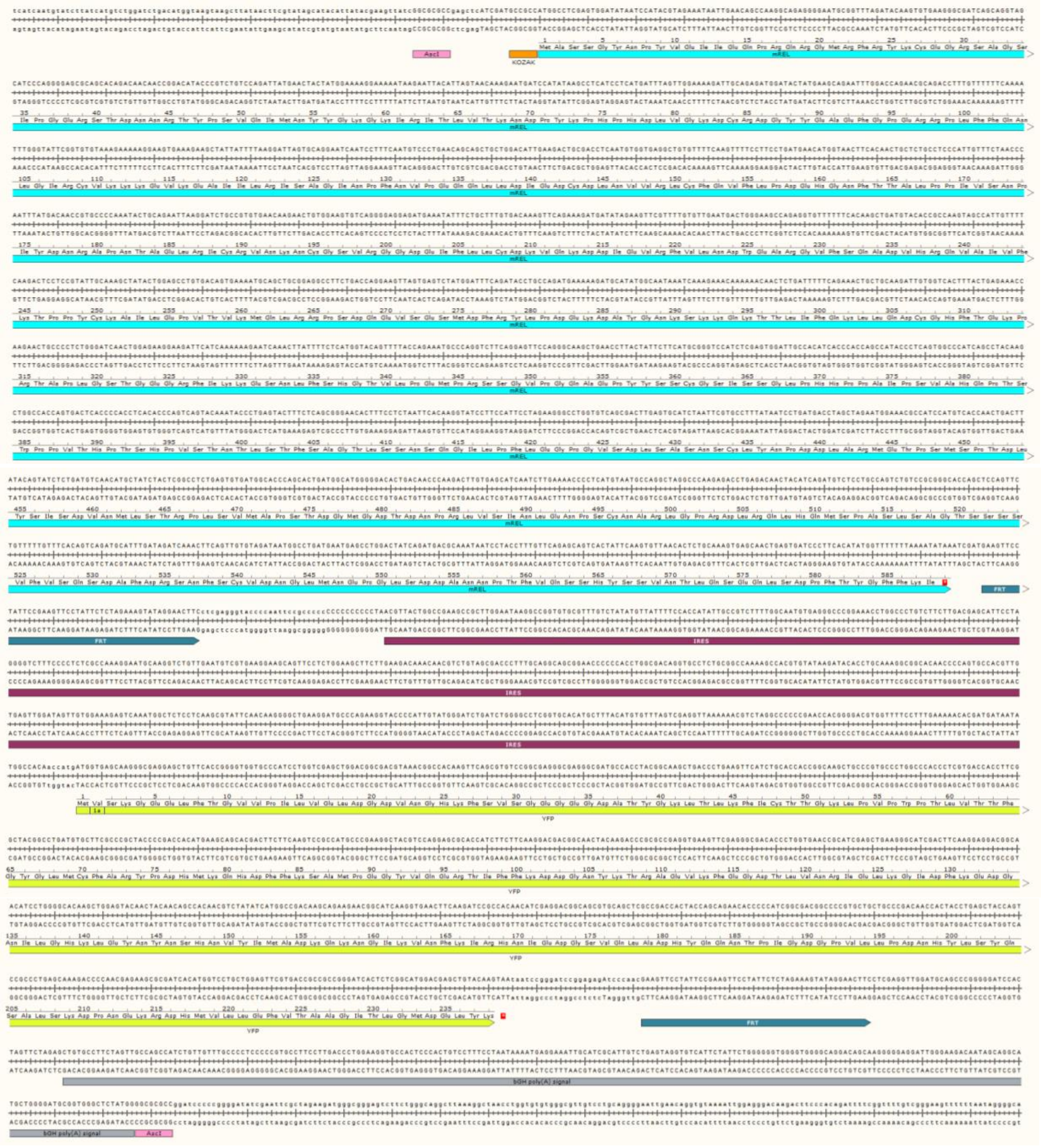


**The sequence of the mREL-YFP insert.** A 3 521 pb AscI/AscI fragment containing the murine REL cDNA (bleu box) followed by an internal ribosome entry sequence (IRES, violet box) and the yellow fluorescent protein (YFP, yellow box) coding sequence, was inserted into the pROSA26-1 vector. The IRES and YFP segment are flanked of FRT sites to removed it if needed. The bGH poly(A) signal and the KOZAK sequence are indicated by a grey box and an orange box respectively. DNA and amino acid sequences are given. Amino acids are numbered.

**PCR screening of ES clones and mice**

Primers sequences for screening REL-YFP ES cells and transgenic mice were 1) for recombined Rosa26 allele, 5pROSA_arm: 5'-CGCCTAAAGAAGAGGCTGTG-3'; neo1: 5'-GGA TGA TCT GGA CGA AGA GC-3', 2) for Rosa26 wild type allele, Rosa_fw: 5'-CTC TCC CAA AGT CGC TCT G-3'; Rosa_rev: 5'-TAC TCC GAG GCG GAT CAC AAG C-3'. Primers sequences for screening AID.Cre$^{ert2}$-TOM model were 1) for tdTomato insertion in Rosa26 locus, tdTOMATO MUT_fw: 5'-CTG TTC CTG TAC GGC ATG G-3'; tdTOMATO MUT_rev: 5'-GGC ATT AAA GCA GCG TAT CC-3', 2) for Rosa26 wild type allele, tdTOMATO WT_fw: 5'-AAG GGA GCT GCA GTG GAG TA-3', tdTOMATO WT_rev: 5'-CCG AAA ATC TGT GGG AAG TC-3', 3) for AID wild type allele, AID_fw: 5'- GTA GGT CCA GCC ATC AGC AG-3'; AID_rev: 5'-CGA AGG TGG CCG AAG TCC AG-3', 4) for Cre insertion into AID locus, Cre_rev: 5'-AGG TTC TGC GGG AAA CCA TTT CCG-3'. Primers sequences for screening CD19-Cre mice were for CD19_Cre allele, 1) for recombined CD19 wild type allele, CD19c: 5'-AAC CAG TCA ACA CCC TTC C-3'; CD19d: 5'-CCA GAC TAG ATA CAG ACC AG-3', 2) for recombined CD19 allele, CD19Cre7: 5'-TCA GCT ACA CCA GAG ACG G-3'.

**Details for generation of AID-TOM mouse model**

To generate the AID-TOM model, we used the AID.Cre$^{ert2}$ and the Ai14 strains. AID.Cre$^{ert2}$ strain is a knock-in transgenic mouse in which a tamoxifen-inducible Cre recombinase enzyme (in fusion with the ERT2 estrogen receptor) is driven by the promoter of the gene encoding the Activation-Induced Cytidine Deaminase AID (Aicda gene, located on the chromosome 6) on one allele. The Ai14 strain is a Cre-reporter mouse with the Rosa26 locus (on the chromosome 6) modified by targeted insertion of a construct containing the strong and ubiquitous CAG promoter, followed by a floxed-stop cassette-driven tandem dimer Tomato (tdTomato) coding sequence on one allele. After breeding homozygous AID.Cre$^{ert2}$ and Ai14 mouse models, F1 double heterozygous mice were obtained and further crossed while always keeping those pups having conserved both independent mutations and until getting animals homozygous for the loss of wild-type AID. As the distance between the AID and Rosa26 loci is close to 10 megabases, which is about 10 centimorgans, we obtained a mouse strain with both transgenes on the same chromosome 6 thanks to a crossing-over recombination event by selecting animals with both the Tomato marker and the AID.Creert2 transgene.

**Housing conditions**

Animals were housed at 21–23°C with a 12-hour light/dark cycle. All procedures were performed under an approved protocol according to European guidelines for animal experimentation (French national authorization number: 8708503 and French Ethics Committee registration number APAFIS#26105-2020061810023698 v1).

**Détails for proliferation experiments**

For proliferation experiments with the Cell Trace Violet, the labeled cells were then seeded at 200,000/well in a 96-well plate with different stimuli: 2.5 µg/mL anti-CD40 monoclonal antibody (HM40-3 clone; eBioscience from Thermo Fisher Scientific, Waltham, MA), 20 ng/mL recombinant murine IL4 (PeproTech from Thermo Fisher); 15 µg/mL AffiniPure F(ab')$_2$ Fragment Goat Anti-Mouse IgG and IgM (H+L) polyclonal antibody (Jackson ImmunoResearch Europe, Cambridge, UK); 50 µg/mL LPS (Merck, Darmstadt, Germany), 2 µg/mL ODN-CpG (ODN-1826; Invivogen, San Diego, CA), and 2 µg/mL ODN-control (ODN-

2136; Invivogen). Cells were cultured in a complete RPMI-1640 (Gibco from Thermo Fisher) medium containing 10% Fetal Calf Serum (Eurobio, Les Ulis, France), 2 mmol/L L-Glutamine, 1 mM sodium pyruvate, 100 IU/mL penicillin, 100 µg/mL streptomycin, and 50 µM βMercaptoethanol (Gibco from Thermo Fisher). After 4 days in a 37°C incubator under 5% $CO_2$, cells were labeled with anti-B220-APC antibody (1/100; BioLegend, San Diego, CA) and Viakrome808 (1 µL per million cells, Beckman Coulter) and then analyzed by flow cytometry using the CytoFLEX LX (Beckman Coulter).

**Ex vivo germinal center B-cell differentiation**

40L-MEF cells were provided by the laboratory of D Kitamura (23). These fibroblasts express the CD40 ligand (CD154) and secrete the cytokine BAFF. The 40L-MEFs are cultured (37°C incubator and 5% $CO_2$) on Thermo Scientific™ Nunc™ cell culture Petri dishes (10 mm) in complete DMEM-Glutamax medium (Gibco): 10% SVF (Gibco), 1 mM sodium pyruvate, 100 IU/mL penicillin and 100 µg/mL streptomycin. At confluence, 40L-MEF cells were treated with 10 µg/mL mitomycin (Sigma-Aldrich from Merck) for 3 hours. Mitomycin-treated 40L MEFs were seeded with 500,000 sorted B cells in 20 mL complete RPMI-1640 medium containing 1 ng/mL recombinant murine IL4 (BioLegend). The medium was replaced after 48 hours.

B cells were analyzed on days 1, 2, 3 and 4. After cell dissociation with Versene (Gibco) and labeling of 40L-MEFs with a PE-coupled anti-CD40L antibody (1/100, BioLegend), B cells were negatively sorted using the EasySep mouse PE kit (StemCell Technologies, Vancouver, Canada). GC B-cell differentiation was monitored by flow cytometry using GC and post-GC B-cells labeling panels. Apoptosis was also assessed by flow cytometry using AnnexinV-AF647 (BioLegend) combined with B-cell labeling using anti-CD19-BV510 and anti-B220-

BV785 antibodies (1/100; BioLegend). Finally, proliferation was assessed after incorporation of 50 µM EdU for 16 hours. EdU-positive cells were labeled with the BaseClick EdU Cell proliferation kit (Sigma-Aldrich from Merck) according to the supplier's recommendations. Prior to flow cytometry analysis, B cells were labeled with anti-CD19 BV510 and anti-B220 BV785 antibodies.

### Antibodies used for flow cytometry analysis

| Target | Antibody clone | Fluorochrome | Dilution | Company |
|---|---|---|---|---|
| Germinal Center B-cell | | | | |
| CD19 | 1D-3 | BUV395 | 1/100 | BD Biosciences |
| CD38 | 90/CD38 | BUV661 | 1/100 | BD Biosciences |
| IgD | 11-26c.2a | BV421 | 1/100 | BioLegend |
| B220/CD45R | RA3-6B2 | BV510 | 1/100 | BD Biosciences |
| CD95 | Jo2 | BV711 | 1/200 | BD Biosciences |
| Gl7 | GL-7 | AF647 | 1/100 | BD Biosciences |
| CD138/Syndecan-1 | 281-2 | APC-Cy7 | 1/100 | BioLegend |
| Activated B-cell | | | | |
| CD19 | 1D-3 | BUV395 | 1/100 | BD Biosciences |
| B220/CD45R | RA3-6B2 | BV510 | 1/100 | BD Biosciences |
| CD86 | GL-1 | BV711 | 1/100 | BD Biosciences |
| IgM | 11/41 | APC | 1/100 | eBioscience |
| CD80 | 16-10A1 | BV421 | 1/100 | BioLegend |
| Post-Germinal Center B-cell | | | | |
| CD19 | 1D-3 | BUV395 | 1/100 | BD Biosciences |
| CD80 | 16-10A1 | BUV661 | 1/100 | BD Biosciences |
| IgD | 11-26c.2a | BV421 | 1/100 | BioLegend |
| B220/CD45R | RA3-6B2 | BV510 | 1/100 | BD Biosciences |
| CD273/PD-L2 | TY25 | BV711 | 1/100 | BD Biosciences |
| IgM | 11/41 | APC | 1/100 | eBioscience |
| IgG1 | RMG1-1 | APC-Cy7 | 1/100 | BioLegend |
| White Blood Cells Populations | | | | |
| CD19 | 1D-3 | BUV395 | 1/100 | BD Biosciences |
| MAC1/CD11b | M1/70 | BV421 | 1/100 | BioLegend |
| B220/CD45R | RA3-6B2 | BV510 | 1/100 | BD Biosciences |
| GR1/Ly6G | 1A8 | BV711 | 1/100 | BioLegend |

| | | | | |
|---|---|---|---|---|
| CD3ε | 145-2C11 | PerCP-Cy5.5 | 1/100 | BioLegend |
| CD4 | GK1.5 | PE/Fire 700 | 1/100 | BioLegend |
| CD335/NKp46 | 29A1.4 | NKp46 APC | 1/100 | BioLegend |
| CD138/Syndecan-1 | 281-2 | APC-R700 | 1/100 | BD Biosciences |
| CD8a | 53-6.7 | APC-Cy7 | 1/100 | BioLegend |
| VIAKROME808 (in each panel) | | Near-IR 750 | 2 µL/$10^6$ cells | Beckman Coulter |
